# Attention biases the appearance of rapidly alternating stimuli

**DOI:** 10.64898/2025.12.21.695868

**Authors:** Arsiak Nabil Ishaq, Erin Goddard

## Abstract

Attentional selection not only increases behavioural performance and neural information for attended stimuli but, in some instances, induces changes in stimulus appearance. While reported appearance changes have mostly concerned the effects of attending to a particular spatial location, here we report a shift in appearance that occurs when attention is directed to one component of a stimulus rapidly alternating at a single spatial location. Our stimuli included two frames of different colors, which alternated at 3.75Hz or 7.5Hz. Participants reported the perceived dominance of one frame’s color in the reference stimulus, by comparison with a target stimulus. We found that when one stimulus frame was more salient than the other, with a greater likelihood of capturing exogenous attention, there was an illusory perception that the frame was physically present for a greater proportion of time. When the frames were matched in salience, a task encouraging participants to endogenously direct their attention to one frame produced a smaller shift in appearance. Our results add to the body of literature demonstrating that attention can change appearance, in this case altering the appearance of attended versus unattended stimuli that are presented at the same spatial location. Furthermore, our results demonstrate that at the ‘temporal transparent’ alternation rate of 7.5Hz, where rapidly alternating stimuli are perceived as two translucent surfaces, attentional selection of one surface can account for the apparent paradox that feature conjunctions are perceived at stimulus alternation rates higher than predicted by the temporal limits of feature binding for other stimuli.

---

Attentional selection of the sensory information allows fast and efficient processing of the most behaviourally relevant information. It is well-established that attention improves behavioural performance related to attended locations and stimulus features (e.g. Carrasco, 2011; Desimone & Duncan, 1995; Purokayastha et al., 2021; Rossi & Paradiso, 1995; Saenz et al., 2003). Consistent with enhanced representations of attended information, attentional focus boosts the stimulus information carried by neural responses (Barnes et al., 2022; Dermody et al., 2025; Goddard et al., 2022; Goddard & Mullen, 2021; Vaziri-Pashkam & Xu, 2017).

There is growing evidence that in addition to selectively enhancing task-relevant sensory processing, attention also changes visual appearance (Carrasco & Barbot, 2019; Chapman et al., 2023). The perceptual effects of covert spatial attention are often in the direction of attention increasing the apparent strength or intensity of the stimulus, including attention increasing perceived stimulus contrast (Carrasco et al., 2004; Liu et al., 2009; Luo et al., 2024), color saturation (Fuller & Carrasco, 2006), flicker rate (Montagna & Carrasco, 2006), speed (Anton-Erxleben et al., 2013; Turatto et al., 2007) and duration (Mattes & Ulrich, 1998; Seifried & Ulrich, 2011). Here we tested whether attention also changes the appearance of ‘temporally transparent’ alternating stimuli.

‘Temporal transparency’ refers to the perception of two simultaneously present, translucent surfaces, that arises when two visual stimuli are rapidly alternated at the same spatial location, at approximately 5 to 20Hz (Holcombe, 2001; Vigano et al., 2015). When the same visual stimuli are alternated at slower rates (below 5Hz) their alternation is perceived as one frame following the other, whereas at very fast alternation rates (>20Hz) observers only see the average of the two frames, and cannot distinguish the properties of the individual frames (Holcombe, 2001; Vigano et al., 2015).

The phenomenon of temporal transparency provides insight into ‘feature binding’: the process by which the brain integrates different attributes of an object, such as color, shape and motion, into a unified perceptual experience. Many early visual cortical neurons respond to multiple feature dimensions (Friedman et al., 2003; Johnson et al., 2008; Lennie & Movshon, 2005), potentially encoding conjunctions of features in the earliest cortical responses to visual input, yet despite this, there is extensive behavioural evidence that additional processing is required for the visual system to solve the ‘binding problem’ (e.g. Ceja et al., 2020; Clifford et al., 2004; Fujisaki & Nishida, 2010). The perception of feature conjunctions within objects, such as the combination of color and orientation, takes longer, and is subject to greater attentional load limitations, compared with perceiving either feature in isolation (Holcombe, 2009; Treisman, 1982, 1996).

An apparent paradox of temporally transparent stimuli is that although the individual stimulus frames alternate much faster than the temporal limit of feature binding in other stimuli, observers are able to accurately judge the conjunction of features within each frame of the stimulus (Holcombe & Cavanagh, 2001; Moradi & Shimojo, 2004; Vigano et al., 2015). For example, Vigano and colleagues (2015) presented stimuli comprising both color and orientation in the same location, pairing orange with a right orientation tilt and blue with a left orientation tilt and vice versa. At temporally transparent alternation rates (7.5 to 15Hz), participants correctly identified color-orientation pairings across the stimulus frames, demonstrating intact feature binding. However, when participants instead judged the coincidence of the same color and orientation pairs when they were spatially separated, they required much slower alternation rates (∼2.5Hz) to perform the task accurately (Holcombe & Cavanagh, 2001; Vigano et al., 2015). These slower rates, corresponding to ∼200ms per frame, are consistent with a slow limit for detecting the coincidence of spatially separate feature pairs (Holcombe, 2009), and seemingly contradict the finding that feature binding is possible at the much higher rates in temporal transparency, when the individual frames are each presented only very briefly (e.g. 16-33ms).

However, there are several results that suggest it is unlikely that feature binding performance in temporal transparency can be attributed to processing occurring within a single frame, and is instead related to the subjective experience of two simultaneously present, translucent surfaces (Clifford et al., 2004; Holcombe, 2009; Holcombe & Cavanagh, 2001; Moradi & Shimojo, 2004; Suzuki & Grabowecky, 2002; Vigano et al., 2014, 2015, 2017). For instance, the alternation rates at which feature binding performance is most accurate is highly correlated with the subjective experience of two translucent surfaces, and feature binding performance in this range of alternation rates can even exceed performance for stimuli that alternate more slowly but evoke a weaker subjective sense of translucency (Vigano et al., 2015). This suggests that in temporal transparency, feature binding does not occur within individual frames, but within the perceptual ‘objects’ corresponding to the persistent transparent surfaces. There are also periodic fluctuations in performance, at ∼8Hz, when participants judge conjunctions for spatially separate features, but not for temporally transparent stimuli, again consistent with qualitatively distinct processes (Nakayama & Motoyoshi, 2019). Accuracy for reporting feature conjunctions increases with overall duration (number of cycles) of the stimulus, consistent with temporal integration of evidence across multiple cycles of the stimulus (Blaževski et al., 2024).

The nature of the persistent surface representations in temporal transparency is unclear, but is has been proposed to be related to the concept of visible persistence (Coltheart, 1980; Moradi & Shimojo, 2004; Vigano et al., 2015). Visible persistence, defined as the visual experience of a stimulus that continues after its physical offset, is thought to be based on neural mechanisms at multiple stages of the visual pathway, including cortically (Coltheart, 1980). The likelihood of visible persistence varies with properties such as stimulus duration, with briefer stimuli more likely than longer stimuli to show persistence (Coltheart, 1980), and to be integrated together into single ‘object’ rather than segregated into distinct temporal events (Dixon & Dilollo, 1994). If similar processes are occurring in temporal transparency, then as stimulus alternation rate increases, and the duration of each stimulus frame decreases, their visible persistence would increase, at the same time as the gap between successive identical frames decreases, both increasing the likelihood that these frames could merge to form an integrated surface or perceptual layer. If both frames of an alternating stimulus are visible simultaneously as continuous surfaces at the same overlapping location, then the perceptual experience of two translucent layers is a natural interpretation.

To model how feature conjunction information could be perceived by participants in temporal transparency, even when individual frames are presented too briefly for a slow binding mechanism, Vigano et al. (2015) proposed that when the stimulus was perceived as two translucent surfaces, observers could direct their attention to one apparent surface at a time. Attentional selection of one surface as an attended object (Desimone & Duncan, 1995; Duncan, 1984) would boost the responsiveness of neurons coding the attended surface’s color and orientation relative to the feature pair of the unattended surface (Engel, 2002; Johnson et al., 2008), which could enable readout of the stimulus feature pairings. If attentional selection of one surface underpins feature binding at these fast alternation rates, we reasoned that this may increase the perceived dominance of the attended surface.

Here, we tested whether manipulating attention alters the appearance of rapidly alternating stimuli, including at ‘temporally transparent’ alternation rates. Attentional selection occurs over time as well as space, including attentional fluctuations entrained to rhythmic stimuli that are task relevant or irrelevant (e.g. see review by Nobre & van Ede, 2018). For an alternating stimulus, attentional selection of one of the two frames could occur either by entrainment in phase with the onsets of the relevant frame, or, as noted by above, by selection the perceptual ‘object’ of the surface (Desimone & Duncan, 1995; Duncan, 1984). The abrupt luminance changes associated with the onset of each frame of an alternating stimulus could repeatedly draw exogenous attention (Franconeri et al., 2005; Remington et al., 1992; Theeuwes, 1995), leading to entrained attentional fluctuations (Nobre & van Ede, 2018). In either case, whether attention either fluctuates rhythmically with the stimulus or if attentional selection occurs at the level of the persistent surface percepts, we expect that when the stimuli are comprised of two frames that differ in salience, the more salient frame will attract exogenous attention to a greater extent than the less salient frame (Spehar & Owens, 2012). By varying the stimuli and the participant’s task, we included conditions where the stimulus surfaces differed in salience (and the likelihood they would exogenously attract attention), or where the task encouraged endogenous attention to be directed to one surface over the other, or where these effects were combined. Our results revealed that across most conditions, the perceived dominance of the attended surface was higher than the unattended surface. When the reference and target stimuli were presented simultaneously, this effect of attention on appearance was present at both temporally transparent alternation rates (7.5Hz) and for stimuli flickering at slower rates (3.75Hz). When the stimuli were instead presented successively, this effect was only observed at a temporally transparent alternation rate (7.5Hz). These results support the hypothesis that attentional selection of one surface could mediate feature binding at fast alternation rates in temporal transparency, and provide a new example of how attention changes appearance.

## Methods

### Participants

28 participants (20 females, 9 males, aged 18-51; M = 22.21, SD = 7.28) took part in Experiment 1; 25 were UNSW Psychology undergraduates and 3 were lab members. Experiment 2 had 33 participants (26 females, 7 males), aged 18 to 53 (M = 22.17, SD = 8.58), with 29 UNSW Psychology undergraduates and 4 lab members. For Experiment 2, data from 7 participants were excluded (n=3 due to incomplete conditions, n=4 due to low performance on 4AFC task in one or more conditions, detailed below), resulting in data from 26 participants in the final analysis. Experiment 3 had 33 participants (20 females, 9 males, 4 preferred not to answer), aged 17 to 29 (M = 20.15, SD= 2.60; 1 preferred not to answer), including 32 UNSW Psychology undergraduates and 1 lab member. For this experiment, 6 participants were excluded in the final analysis (n=3 due to incomplete conditions, n=3 due to low performance on the 4AFC task in one or more blocks, detailed below), with data from the remaining 27 participants included. An analysis of power suggested that a sample size of n=27 was required to detect a medium effect (Cohen’s *d*=0.5) with power of 0.8, at α error probability of 0.05 (calculated with G*Power v 3.1).

All participants had normal or corrected-to-normal visual acuity, and no history of seizures or epilepsy. Each participant had normal color vision, confirmed by screening with the Hardy Rand Rittler pseudoisochromatic plates (4^th^ edition, published by Richmond Products). The study was conducted according to the protocols approved by the Human Research Ethics Approval Panel – C (HREAPC) within the School of Psychology, UNSW (Reference iRECS4177). The data and materials for both experiments are available at (*URL with DOI will be made available if accepted for publication*).

### Experimental setup and display

Stimuli were presented on calibrated 32-inch Display++ LCD Monitors by Cambridge Research Systems (resolution of 1920 x 1080), using Windows (Version 10) and MATLAB (R2022b), together with routines from Psychtoolbox 3 (Brainard, 1997; Kleiner et al., 2007; Pelli, 1997). Participants completed the experiments in a darkened room, viewing the screen from approximately 60 cm. All stimuli were displayed on a background on mean grey (x=0.293, y=0.326, Y=60 cd/m^2^).

### Color properties and determining subjective isoluminance

All stimuli in Experiments 1, 2 and 3 alternated between ‘orange-blue, or ‘green-magenta’: these color pairs include equal modulations of color contrast along the L-M and S-cone isolating directions of the DKL space (Derrington et al., 1984), but with opposite pairings (as in Goddard et al., 2010). Every stimulus frame was made up of pixels of a single hue at one of two achromatic offsets (a 10% increment and a 10% decrement, relative to the background).

At the start of the experimental session, participants completed a ‘minimum flicker’ task, where we used heterochromatic flicker photometry (He et al., 2020; Smith & Pokorny, 1975) to equate all luminance increment stimuli for subjective isoluminance, and similarly to equate all luminance decrement stimuli. This ensured that in the main experiment, the alternations between colors were equated across participants in their luminance content, avoiding luminance artifacts that could result from differences in each participant’s point of isoluminance. On each trial, participants saw a vertical squarewave grating (0.5 cycles/degree) in a circular aperture (diameter 16.5 degrees) which modulated between mean grey and either an achromatic offset (10% increment or 10% decrement) or a uniform color. The grating stimuli alternated at 10Hz between the achromatic and chromatic components at 10Hz. The achromatic component remained fixed, while participants adjusted the luminance of the chromatic component via keypresses until they minimised the experience of flicker. Participants repeated this process three times for each of the four hues at two luminance offsets, completing a total of 24 trials, which were randomly interleaved. Settings across the three repeats were averaged in each case to determine the 8 unique colors that were used in the experimental stimuli for each participant.

### Experimental stimuli

All experimental stimuli comprised two frames alternating at 7.5Hz, within the temporally transparent range reported by previous studies (Holcombe, 2001; Vigano et al., 2015) (Experiments 1, 2 and 3) or at 3.75Hz (Experiment 2 and 3 only), which evokes a much weaker sense of transparency (Vigano et al., 2015). Each stimulus was 17.76 x 17.76 degrees and alternated over time between two different colored frames. Example stimulus frames are shown in **Figure 1**. In all experiments, participants compared a target stimulus with a reference stimulus, presented either simultaneously on either side of a central fixation marker (Experiments 1 and 2), or successively, at the centre of the screen (Experiment 3), as seen in **Figure 2**.

**Figure 1.**
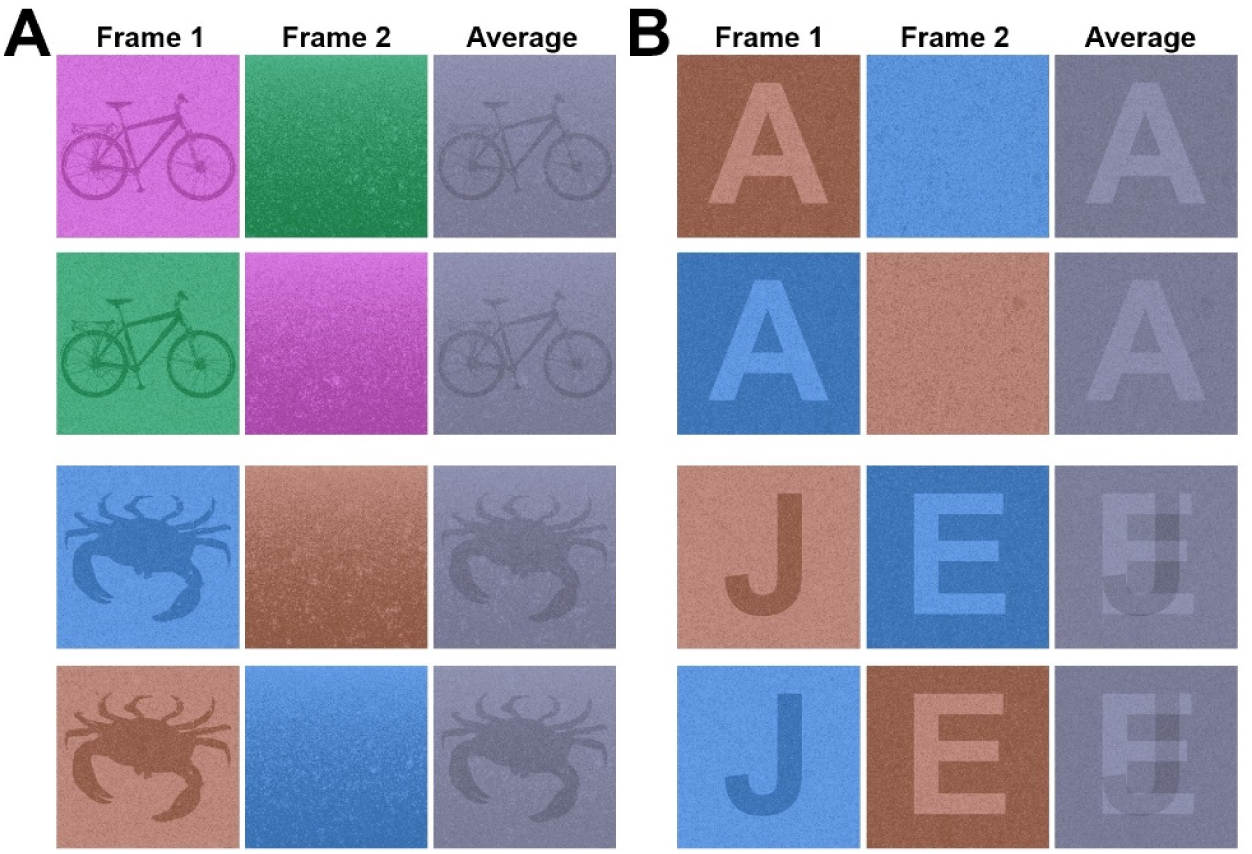
Example stimuli, composed of two frames alternating over time. In each of the 8 examples, the average of the two frames illustrates the fused percept that would result if alternation rates were too fast to perceive the individual frames. The 8 example stimuli yield 4 unique averages, demonstrating the disappearance of feature conjunction information in the fused percept. **A)** Example stimulus frames from Experiment 1: each stimulus included an object silhouette frame and a gradient frame, paired with either magenta and green or orange and blue. **B)** Example stimulus frames from Experiments 2-3: a letter silhouette was either paired with a plain frame or a second letter silhouette, all in orange and cyan.

**Figure 2.**
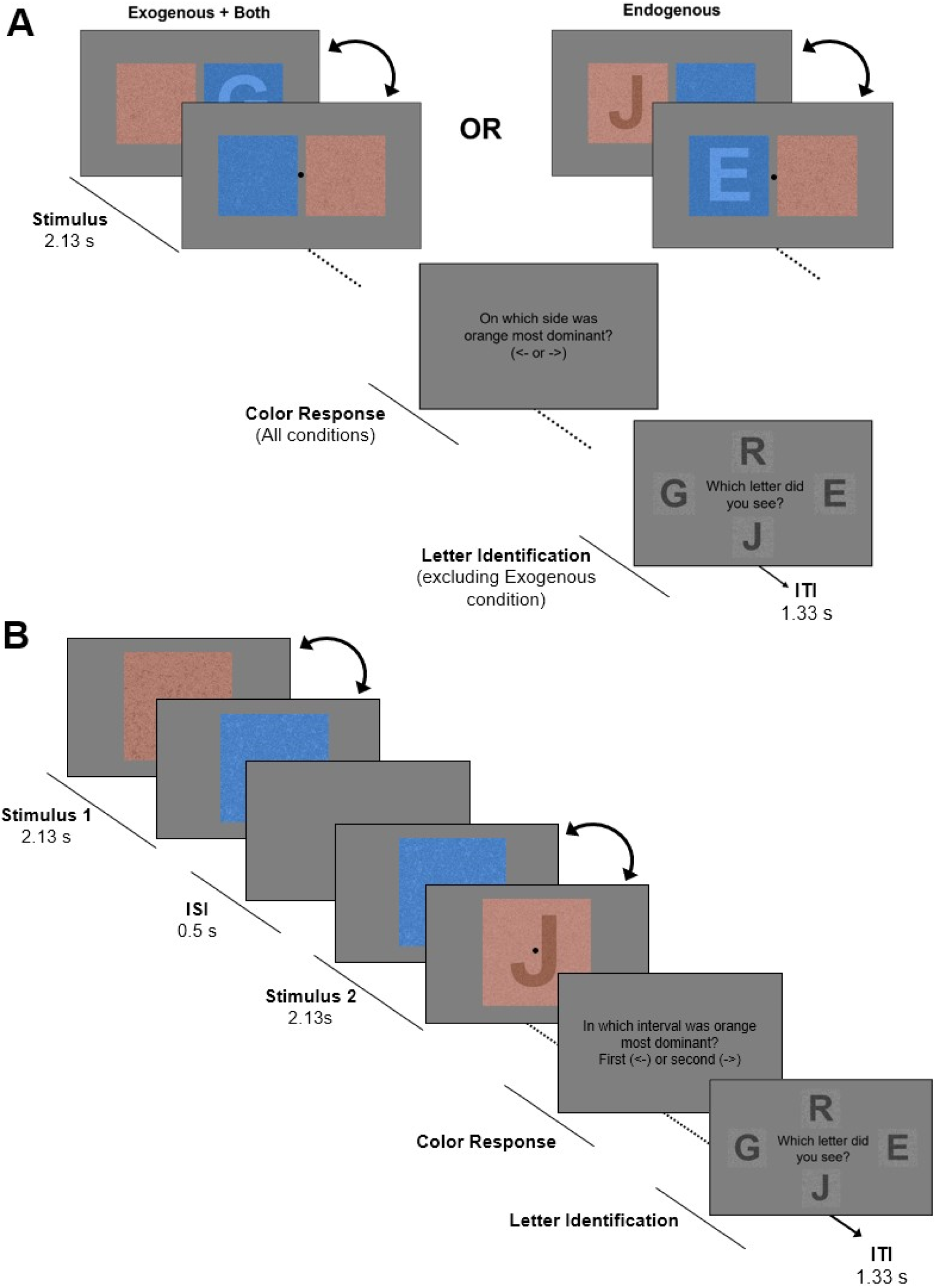
Trial timelines for Experiments 1-2 (**A**) and Experiment 3 (**B**). **A)** In Experiments 1 and 2, all trials commenced with two alternating stimuli (a target and a reference stimulus) to the left and right of a central fixation marker, followed by a judgement of color dominance. Attention conditions were either ‘Exogenous’ (where one stimulus frame was more salient than the other other), ‘Endogenous’ (where the object/letter silhouette was task relevant) or ‘Both’, see text for details. In the ‘Endogenous’ and ‘Both’ conditions, participants then identified the object or letter they saw from amongst distractor items. **B)** In Experiment 3, the two stimuli to be compared were presented in two intervals rather than side-by-side, remaining details were the same as in Experiments 1-2. **ITI** = inter- trial interval. **ISI** = inter-stimulus interval. An instruction screen was shown at the start of each block, and a blank sen with a fixation marker was shown during the **ITI** (not shown in timeline).

#### Experiment 1

In Experiment 1, the reference stimuli included an object ‘silhouette’ frame and a simple gradient frame (example shown in **Figure 1A**), while the target stimuli comprised two gradient frames. The object images used in Experiment 1 were sourced from a previous study by Contini (2020): 32 (of the original 120) stimulus images were converted to silhouettes, in each case as a darker object on a lighter background. A uniform noise filter was applied to each silhouette image, where 1/6 pixels were randomly selected for polarity inversion. The silhouette frames alternated with a noise texture with a linear gradient, where pixels varied from 92% light to 92% dark. For half of the reference stimuli, the gradient image was vertically flipped. All target stimuli in Experiment 1 included two gradient frames, one of each direction.

#### Experiment 2

In Experiment 2, we included a condition where participants attended to just one of the frames in the reference stimulus (while ignoring a stimulus in the alternate frame). Since the objects used in Experiment 1 varied in their similarity to one another, and in their mutual interference when superimposed, for Experiment 2 we switched to reference stimuli that included a letter ‘silhouette’ frame, which was either paired with a second letter silhouette, or a frame of plain color (example shown in **Figure 1B**). The letters were one of eight uppercase letters (A, E, G, J, M, Q, R, or T) converted into a plain silhouette. The set of letters were selected to minimise overlapping features and reduce confusability when superimposed (e.g. avoiding pairs such as ‘P’ and ‘R’, where ‘P’ is a subset of the ‘R’ shape). The letter silhouette frames were either a dark letter on a lighter background, or a light letter on a darker background. When the reference stimulus included two frames with letter silhouettes, there was one of each luminance polarity. When the reference stimulus had a letter alternating with a plain frame, the plain frame was of the opposite luminance polarity to the background of the letter frame. In both cases, the corresponding target stimulus included two frames of the same color and luminance as the letter background(s) or the plain frame. As in Experiment 1, a noise filter was applied to all silhouette stimuli, with 1/6 of pixels selected for polarity (light/dark) inversion. For Experiment 2, the same filter was applied to the plain frames.

#### Experiment 3

Experiment 3 used the same letter stimuli as in Experiment 2, but the reference stimulus always included a single letter (as in the upper example of **Figure 1B**).

### Experimental design

All experiments employed a within-subjects, method of constant stimuli design. We measured any shift in the perceived dominance of the attended frame in the reference stimulus by varying the relative duration of the two frames in the target stimulus. In Experiments 1 and 2, we asked participants to report on which side (left or right) a pre-specified color (e.g. orange) appeared more dominant (two alternate forced choice, or 2AFC, design, **Figure 2A**). In Experiment 3, participants instead reported the interval (first or second) in which the color appeared more dominant (two interval forced choice, or 2IFC, design, **Figure 2B**). Participants were informed of the color they were to report at the beginning of each block of trials and instructed to keep their eyes on the central fixation marker (a small black circle) at the start of each trial and during the stimuli, and to covertly attend to the stimuli presented on the left and right (Experiments 1-2) or centrally (Experiment 3). On our 120Hz displays, the 7.5Hz stimulus was 16 frames per cycle. In the reference stimulus, both frames were always of equal duration (8 frames each per cycle), whereas in the target stimulus, the 16 frame cycle included either 3, 5, 6, 7, 8, 9, 10, 11 or 13 frames of one color, with the remaining frames of the alternate frame; trials of different target stimulus relative durations were randomly interleaved within each block of trials. In the 3.75Hz condition (Experiment 2 and 3), the cycle was twice the duration, but the same set of relative frame durations were used. Target and reference stimuli were either presented simultaneously (Experiments 1 and 2) or successively (Experiment 3 only) with randomly offset starting phases, meaning that both stimuli could commence the trial at any point in their cycle, and the reference stimulus was equally likely to start with either frame. The locations (left or right of fixation in Experiments 1 and 2; before or after fixation in Experiment 3) were counterbalanced across trials. Finally, the color of the object or letter in the reference stimulus was counterbalanced across trials in each block. This meant that even when there is a strong shift in perceived dominance of the reference stimulus, participants would always give approximately even numbers of responses ‘left’ and ‘right’, or ‘first’ and ‘second’, and responses selecting target and reference stimuli (e.g. if reporting the side on which orange is more dominant, a shift in perceived dominance would lead to more reference side responses when the attended reference frame is orange, but more target side responses when the attended reference frame is blue).

#### Experiment 1

In Experiment 1, stimuli were presented for 1.07 s, followed by the 2AFC color response. To encourage participants to endogenously direct their attention to the object silhouette, on each trial the color response was followed by a four alternate forced choice (4AFC) judgement of which object silhouette they saw. The 4AFC included a greyscale version of the silhouette they saw, along with 3 other objects randomly selected from the set of 32 objects, similar to the last frame of **Figure 2A**. Participants indicated their response with an arrow key. Feedback was provided following the completion of each block regarding their accuracy on the 4AFC object identification task; however, no feedback was given on the color task. Each participant completed four blocks of 88 trials (judging the dominance of blue, orange, green and magenta across different blocks), totalling 352 trials. Block order was counterbalanced across participants.

#### Experiment 2

In Experiment 2, we tested how flicker rate and attention type affect stimulus appearance. Experiment 2 included a similar design to Experiment 1, but with a reduced number of color conditions (all stimuli were orange-blue), and two stimulus alternation rates (7.5Hz or 3.75Hz), in three attention conditions (exogenous only, endogenous only and both). We manipulated attention condition by varying the stimuli and secondary tasks. The ‘both’ condition was equivalent to Experiment 1: the reference stimulus had a letter in just one frame (increasing its salience and the likelihood of attracting exogenous attention to this frame), and each trial included a 4AFC task where participants identified the letter they saw (encouraging participants to endogenously direct their attention to the letter). In the ‘exogenous only’ condition, the stimulus remained the same, but participants were told they could ignore the letter, and the 4AFC task was removed. In the ‘endogenous only’ condition, we equated the salience of both frames of the reference stimulus by adding a task-irrelevant letter to the alternate frame, to equate these frames in their likelihood of exogenously attracting attention. Participants attended either to the light letters on dark backgrounds, or the dark letters on light backgrounds (this instruction was consistent across blocks for each participant but counterbalanced across participants). In the ‘endogenous only’ condition, the letter options in the 4AFC task included both the task-relevant and the distractor letters, along with 2 other letters randomly selected from the set of 8 letters. Across all conditions, stimuli were presented for 2.13 s before participants were prompted to make a color response. This duration was doubled (relative to Experiment 1) to account for the increased task difficulty in the ‘endogenous’ condition. In the ‘both’ and ‘endogenous only’ attention conditions, participants were given feedback on their letter identification accuracy at the end of each block. Each participant completed one block of 56 trials for each combination of alternation rate (7.5Hz or 3.75Hz), color reported (blue or orange) and attention condition, totalling 12 blocks and 672 trials. The order of attention conditions, alternation rates and color reporting (blue or orange) were counterbalanced across participants.

#### Experiment 3

In Experiments 1-2, although we sought to test the effect of attending to one of two frames in each stimulus, our design included a spatial comparison of two stimuli presented simultaneously, making it likely that participants covertly directed their attention to each side in turn to make the comparison. To test whether our findings generalise to a design excluding this component, where there is no difference in spatial attention between the target and reference stimuli, we conducted a third experiment. In Experiment 3, we employed a similar design to Experiment 2, except that stimuli were presented successively instead of simultaneously to limit spatial effects. Both stimulus alternation rates (7.5Hz or 3.75Hz) were included, however, only the ‘both’ condition of Experiment 2 was tested. This included reporting which interval a color was more dominant in (two interval forced choice, 2IFC) and which letter was present (4AFC). Each interval was presented for 2.13 seconds, with a gap of 0.5 seconds. Participants each completed four blocks of 56 trials, with the order of alternation rates and color reporting counterbalanced across all participants.

### Analysis

To exclude data from participants who may not have been following task instructions, we excluded all data from any participant with outlier accuracy, defined as lower than 2 quartiles below the mean in any of the 4AFC tasks (n=4 excluded from Experiment 2, n=3 excluded from Experiment 3). To estimate the perceived dominance of the attended frame in the reference stimulus, we calculated the point at which the target stimulus was perceived to have the same proportion of the two colors as the reference stimulus (point of subjective equality, PSE). To do this, we constructed a series of psychometric functions, defined as the likelihood that the color of attended stimulus appeared more dominant in the target (*F*(*x*; α, *β*)), as a function of the proportion of that color in the target stimulus (*x*). Data were fit with a Weibull psychometric function:

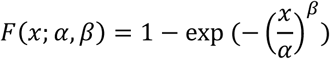

where α is the PSE, and *β* is a the slope of the function (Kingdom & Prins, 2016). In each case, the MATLAB function *fminunc* was used to find the best fitting values for PSE and slope using a gradient descent approach. We fit these psychometric functions to data from each individual participant, for each attention condition (Experiment 2 only), and alternation rates (Experiments 2 and 3) collapsing across location of the target stimulus (left or right, or first or second) and the color whose dominance the participant was reporting.

### Statistical Testing

In Experiment 1, we used a non-parametric permutation test (Efron & Tibshirani, 1994) of whether the PSE values were significantly biased away from 0.5, equivalent to a two-way t-test. We generated 10,000 bootstrapped samples of the PSE data, in each case sampling the PSEs from the 28 participants a total of 28 times, with replacement. To implement a two-way test of whether the observed PSE was different from 0.5, we calculated the p-value as twice the smaller of two proportions: the sample above and below 0.5.

In Experiment 2, we tested whether there was evidence of a significant bias in the average PSE of each condition using a series of two-sided permutation tests equivalent to that used in Experiment 1, with a false discovery rate (FDR) correction for multiple comparisons (Benjamini & Hochberg, 1995). To test whether the magnitude of the shifts in PSE varied across conditions, we conducted a 2-way repeated measures ANOVA, testing the effects of attention condition and stimulus flicker rate on PSE. To estimate the p-values for these statistics non-parametrically, we performed the same ANOVA on a series of randomly permuted datasets, that were generated by shuffling the condition labels for the PSE data using a separate random seed for each participant’s data. We repeated this permutation process 10,000 times, generating a null distribution of each F statistic to which we could compare our observed values. In each case, the p-value was calculated as the proportion of F statistics in the null distribution that were greater than the observed value. Similarly, for follow up pairwise contrasts, we compared the same null distribution to with the observed differences in each case, then applied an FDR correction for multiple comparisons (Benjamini & Hochberg, 1995).

In Experiment 3, we tested whether there was evidence of a significant bias in the average PSE of each condition using two two-sided permutation tests, equivalent to those in Experiment 1, with an FDR correction for multiple comparisons. To test for evidence of a significant difference in PSEs between the two conditions we used a permutation test equivalent to a paired t-test, using 10,000 permutations to generate a null distribution of the difference between the two conditions, to which we compared the observed value.

## Results

### Experiment 1

In Experiment 1, participants completed two tasks on each trial, judging whether a particular color was more dominant in the stimulus to the left or to the right of fixation, and reporting which object silhouette they saw in the reference stimulus. All participants achieved a high level of accuracy on the object identification task (average 96.3% correct, 95% CI [95.3-97.3%], where chance rate is 25%), and no participants were excluded as outliers. Note that performance on this task does not provide evidence of an attentional boost in performance, but suggests that participants were following instructions for the task that encouraged them to direct their endogenous attention to the object silhouette during the reference stimulus.

Results of the color dominance task are shown in **Figure 3**. If there were no effect of attention on the apparent dominance of the attended color, then the psychometric curve in **Figure 3A** would go through the middle of this plot, where the red dashed lines intersect at 0.5. Instead, the data in **Figure 3A** is shifted towards the right, meaning the target stimulus tended to appear equivalent to the reference stimulus when the target stimulus was biased towards the attended color. This trend was consistent across data from individual participants, as seen in the distribution of PSEs in **Figure 3B**. A non-parametric two-way test showed the observed average PSE of 0.558 (95% CI [0.546 0.570]) was significantly greater than 0.5 (p<0.001).

**Figure 3.**
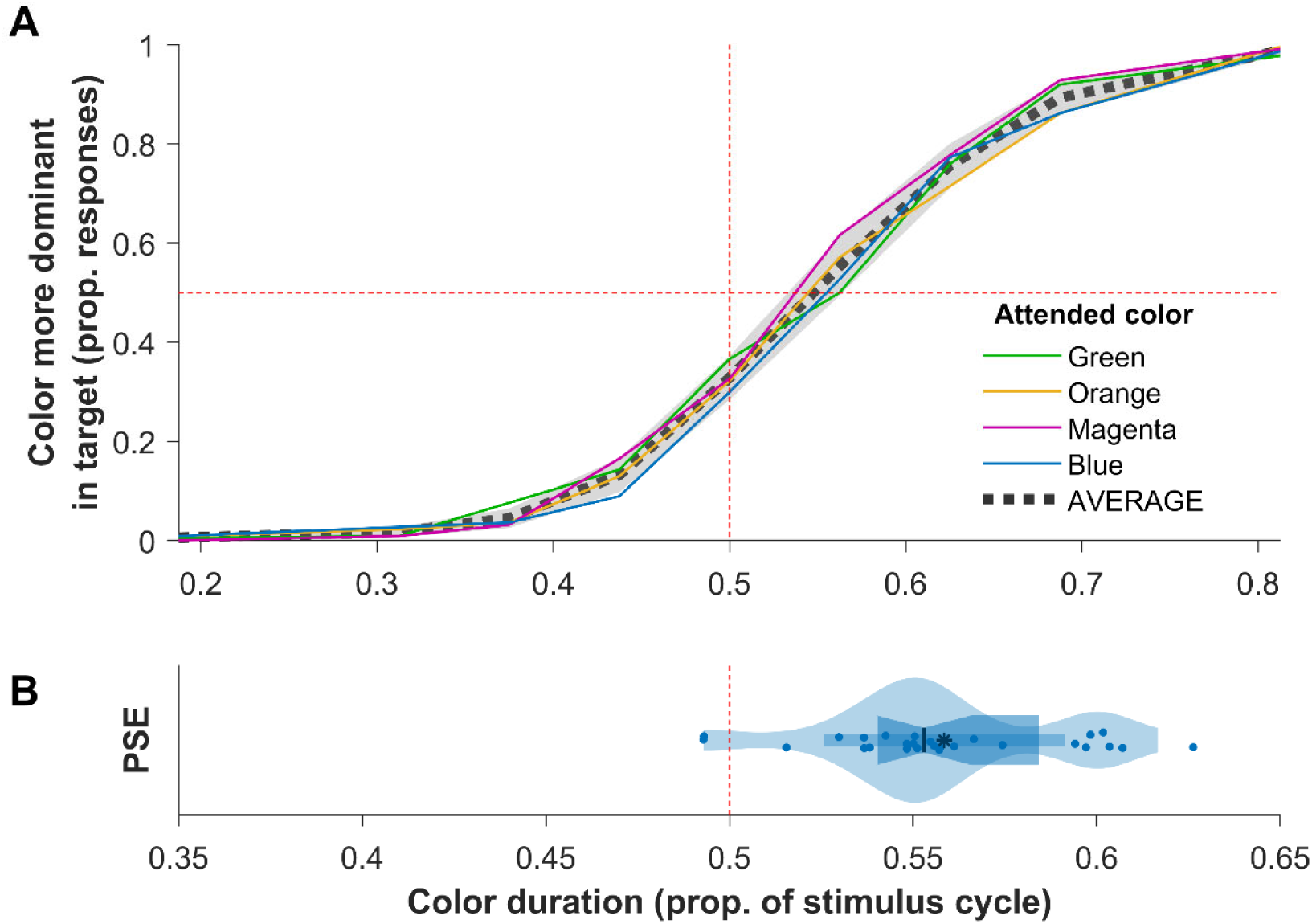
Group data (n = 28) showing the shift in the perceived dominance of the attended color (the color of the object silhouette in the reference stimulus) in Experiment 1. **A)** The proportion of responses where the reference stimulus was judged more dominant in the attended color. The colored lines show the average responses for each attended color. Average data (black dashed line) are shown with shaded error bars indicating 95 % CIs of the between-subject mean. **B)** Violin plot showing the distribution of points of subjective equality (PSEs, dots show individual participants), obtained when each participant’s average data were fit with a psychometric function (see text for details, fits not shown).

These data suggest that the reference stimulus, in which the alternating frames were of equal duration, appeared biased towards the color of the frame containing the object silhouette. That is, attending to the frame with the object silhouette changed the overall appearance of the stimulus, increasing the apparent dominance of the attended frame and/or reducing the dominance of the alternate frame.

### Experiment 2

In Experiment 2, we replicated the findings of Experiment 1 with a new cohort of participants, and a modified stimulus (using letter silhouettes instead of objects). In addition, we extended our original design in two ways: first, we tested whether this effect is specific to stimuli in the temporally transparent range, by including stimuli that alternated at 3.75Hz. Second, we included three different attention conditions, in order to test the extent to which the shift in appearance was dependent on the task-relevance of the stimulus, which would suggest it may be driven by the endogenous direction of attention, or task independent, suggesting it is driven by the salience difference between frames in the reference stimulus. After excluding 4 participants with low accuracy in the 4AFC task in one or more conditions (lower than 2 quartiles below the mean), remaining participants achieved a high level of accuracy on the object identification tasks, with high performance in the ‘both’ attention condition (7.5Hz: average 98.1% correct, 95% CI [97.3-98.9%]; 3.75Hz: average 96.4% correct, 95% CI [95.4-97.3%]), and the ‘endogenous only’ attention condition (7.5Hz: average 98.3% correct, 95% CI [97.7-98.9%]; 3.75Hz: average 94.5% correct, 95% CI [92.8- 96.3%]), in each case well above chance rate (25%).

The shifts in appearance for both flicker rates across each attention condition are shown in **Figure 4**. As in Experiment 1, the PSEs tended to be above 0.5, indicating that attention increased the apparent dominance of the attended frame in the reference stimulus. A series of permutation tests (see methods for details) showed that the average PSE was significantly biased (significantly greater than 0.5) across five of six conditions (p<0.001 in all cases except in the endogenous only attention condition for the 7.5Hz stimuli, where the p-value did not survive FDR correction at q<0.05). A repeated measures ANOVA with bootstrapped p-values revealed significant main effects of attention condition (F(2,50)=25.66, p<0.001), and stimulus flicker rate (F(1,25)=6.69, p=0.013), but no significant interaction between these factors (F(2,50)=2.03, p=0.14). That is, at 3.75Hz, the reference stimuli appeared significantly more biased towards the color of the attended frame than for the temporally transparent stimuli at 7.5Hz.

**Figure 4.**
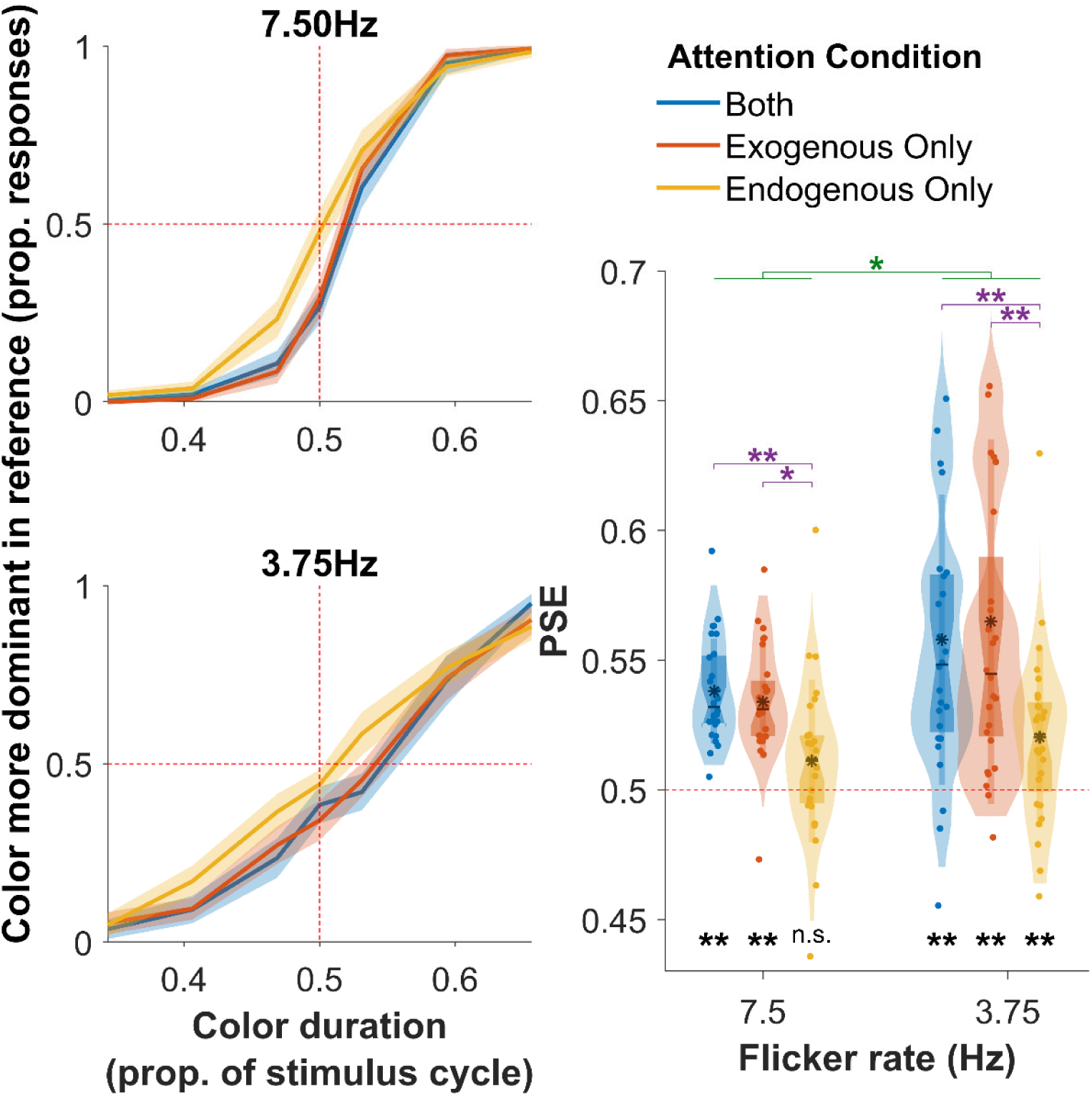
Group data (n=26) showing shifts in appearance across stimulus flicker rates and attention conditions (Experiment 2). Average responses across conditions (left) show shifts in perceived color dominance at 7.5Hz (top) and 3.75Hz (bottom). Shaded error bars indicate 95% CIs of the between- subject mean. Violin plots (right) display the distributions of PSEs (dots show individual participants), for the three attention conditions and two flicker rates. Black asterisks indicate where the relevant data were significantly different from 0.5; purple and green asterisks indicate there were significant differences between PSEs across conditions (* p<0.05; ** p<0.01, with FDR correction at q<0.05).

To further investigate the effect of attention, we conducted pairwise contrasts between the attention conditions, within each stimulus flicker rate. These revealed a significantly more biased PSE in the ‘both’ condition than in the ‘endogenous only’ condition, for both the 7.5Hz stimuli (p=0.008) and the 3.75Hz stimuli (p<0.001). The average PSE in the ‘exogenous only’ condition was significantly greater than in the ‘endogenous only’ condition for both the 7.5Hz stimuli (p=0.031) and the 3.75Hz stimuli (p<0.001). The ‘exogenous only’ attention condition was not significantly different from the ‘both’ condition for either the 7.5Hz or 3.75Hz stimuli (p-values did not survive FDR correction at q<0.05).

We note that the slopes of the psychometric functions at the slower alternation rate tended to be shallower (**Figure 4**), suggesting that participants were less precise in judging the relative frame durations. This difference in slope is present even when the data were expressed as a proportion of the stimulus cycle: in frame duration the differences in slope are even greater. Our design was not optimised for estimating the slope of these curves, which require substantially more data than for estimating points of subjective equality (Kontsevich & Tyler, 1999), but this could be investigated in future work.

### Experiment 3

In Experiment 3, we replicated the findings from Experiments 1 and 2 with a new cohort for stimuli presented successively in two intervals, rather than side by side. In the two interval design of Experiment 3 we excluded any effects that could be related to differences in spatial attention between the two stimuli within each trial in Experiments 1 and 2. Only 3 participants were excluded to low performance on the 4AFC task (below 0.85, >2 quartiles below mean). Remaining participants achieved a high level of accuracy on the letter identification task (7.5Hz: average 96.2% correct, 95% CI [95.3-97.2%]; 3.75Hz: average 96.5% correct, 95% CI [95.5-97.5%).

The shifts in appearance for both stimulus alternation rates are shown below in **Figure 5**. At the temporally transparent alternation rate (7.5Hz), the PSE was significantly greater than 0.5 (p=0.016), consistent with the color of the attended surface appearing more dominant than the gradient surface in the reference stimulus, as in Experiments 1 and 2. At the slower alternation rate, the PSE was not significantly different from 0.5 (p-value was not significant after FDR correction at q<0.05), and was significantly lower than the PSE in the 7.5Hz condition (p<0.001, two-tailed permutation test).

**Figure 5.**
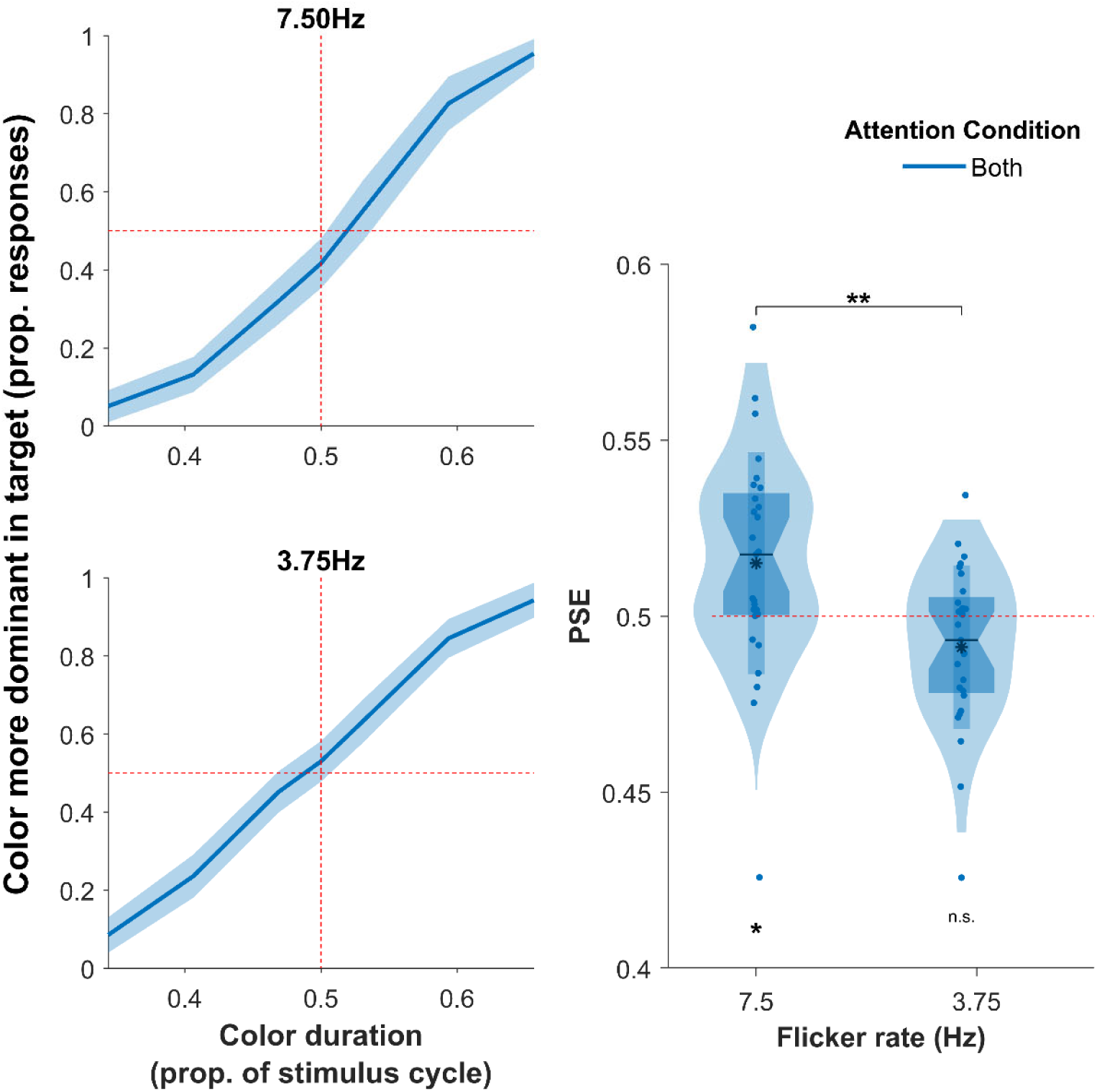
Group data (n=27) showing shifts in appearance across stimulus flicker rates (Experiment 3). Average responses (left) showing shifts in perceived color dominance at 7.5Hz (top) and 3.75Hz (bottom). Shaded error bars indicate 95% CIs of the between-subject mean. Violin plots (right) display the distributions of PSEs (dots show individual participants), for the two flicker rates. Black asterisks indicate where the relevant data were significantly different from 0.5; purple and green asterisks indicate there were significant differences between PSEs across conditions (* p<0.05; ** p<0.01, with FDR correction at q<0.05).

## Discussion

In this study, we sought to test whether attending to one of two frames in an alternating stimulus changes the appearance of the stimulus, making the attended frame seem more dominant, and perceived as if it were physically present for a greater proportion of the stimulus cycle. Across multiple stimuli and attention conditions, we found evidence that attention produces these shifts in appearance. For stimuli alternating at 7.5Hz, consistent with the ‘temporally transparent’ percept of two simultaneously present, overlapping surfaces (Holcombe & Cavanagh, 2001; Vigano et al., 2015), we found the more salient surface appeared more dominant (Experiments 1-3), with a weaker effect in the same direction that did not reach statistical significance when the stimulus frames were matched in salience but the task encouraged participants to endogenously direct their attention to one frame (Experiment 2). At the slower 3.75Hz alternation rate, where there is little or no percept of temporal transparency, we found effects in the same direction when participants judged stimuli that were presented side by side (Experiment 2), but we did not find any evidence of this effect when the two stimuli were presented sequentially (Experiment 3, where there was a non-significant trend for an effect in the opposite direction).

Where the stimulus frames of the alternating stimulus differed in salience (in Experiments 1 and 3, and the ‘exogenous only’ and ‘both’ conditions of Experiment 2), we reasoned that each onset of the more salient stimulus could exogenously draw attention (Remington et al., 1992; Theeuwes, 1995) to a greater extent than the onsets of the less salient alternate frames (Spehar & Owens, 2012).

Salience can also modulate visible persistence, or the perceptual experience of a stimulus continuing after its physical offset, but in the opposite direction: ‘the inverse intensity effect’ describes the tendency for more intense stimuli to have briefer persistence (Coltheart, 1980), so we consider the attraction of exogenous attention as the most likely account of our findings.

There are two main implications of these findings: they add to the body of work demonstrating ways in which attention biases the appearance of the attended stimulus (Carrasco & Barbot, 2019). Second, this work supports the hypothesis that feature binding for temporally transparent stimuli could be mediated by attentional selection of the apparent ‘surfaces’ (Vigano et al., 2015).

### Attention changes the appearance of alternating stimuli

Our finding that attention increased the perceived dominance of the attended frame is consistent with attention increasing the ‘strength’ of the percept, including aspects like contrast and color saturation and speed (Carrasco et al., 2004; Fuller & Carrasco, 2006; Liu et al., 2009; Luo et al., 2024; Turatto et al., 2007). For briefly presented (rather than alternating) stimuli, directing attention to the spatial location of a stimulus increases its perceived duration relative to when attention is directed to other locations (Enns et al., 1999; Mattes & Ulrich, 1998; Seifried & Ulrich, 2011), or another sensory modality (Mattes & Ulrich, 1998). However, our finding that attention increases the perceived dominance of the attended frame cannot be accounted for by these previously reported effects, since in our case the unattended frames, which were perceived as relatively weaker or shorter in duration, were at the same spatial location as the attended frame, rather than in an unattended location. This means that our findings cannot be attributed to any effects on appearance that are driven by attending to a particular spatial location alone. The attentional effects we observe may suggest that during the alternating stimuli, participants varied their attentional focus on the stimulus in sync with the alternation rate, modulating between stronger and weaker attentional engagement with the stimulus as its frames updated, as rhythmic entrainment with the stimulus (Nobre & van Ede, 2018). In this scenario, stronger attentional engagement in the attended frames could increase their perceived duration, similar to that of briefly presented stimuli (Enns et al., 1999; Mattes & Ulrich, 1998; Seifried & Ulrich, 2011). Consistent with this idea, performance-enhancement from attention includes rhythmic components that may be linked to neural oscillations, especially in the theta (∼4-8Hz) and alpha (∼8-12Hz) range (Shalev et al., 2019). Behavioural evidence suggests that spatial and object-based attention, for a non-rhythmic stimulus, fluctuate over time at 4-8Hz (Fiebelkorn et al., 2013; Landau & Fries, 2012), which is in approximately the same range as the frequency of our stimuli (3.75 and 7.5Hz), although note that for rhythmically alternating stimuli, these oscillations were found only for spatially separated stimuli, and not for spatially superimposed stimuli (Nakayama & Motoyoshi, 2019). To mediate the attentional effects observed here would require that these modulations flexibly become phase-aligned with the appropriate stimulus: supporting this, rhythmically presented visual stimuli can entrain neural oscillations (Calderone et al., 2014; Lakatos et al., 2008; Mathewson et al., 2012), which could mediate the effects we find of exogenous attention changing appearance. In the ‘endogenous only’ condition, where we found weaker but significant effects of attention on appearance, syncing of attention to the stimulus phase cannot be accounted for by stimulus-driven entrainment alone. However, predictable, but non- rhythmic, stimuli of a similar alternation rate can produce a distinct yet similar attentional enhancement to rhythm-based effects (Breska & Deouell, 2017), demonstrating that memory-based predictions can also modulate attentional fluctuations. Our finding of an endogenous attention effect suggests a similarly internally driven control of these attentional fluctuations.

An alternative account of our results, which is not mutually exclusive with the idea of attentional fluctuations, is that they are mediated by attentional selection of one of the persistent surfaces that are perceived in temporal transparency (Holcombe & Cavanagh, 2001; Vigano et al., 2015). Object- based attention accounts for the selection of one surface, even where spatially superimposed with other objects or surfaces (Desimone & Duncan, 1995; Duncan, 1984). In our task, when participants attended to a task-relevant object silhouette, or letter (or if their attention was exogenously captured by the task-irrelevant but salient letter), an object-based attention account posits that all features of the attended surface would be selected (Duncan et al., 1997), which could increase the intensity or dominance of the attended surface’s color, in a similar way to spatial attention’s effects of increasing perceived stimulus contrast (Carrasco et al., 2004; Liu et al., 2009; Luo et al., 2024), and color saturation (Fuller & Carrasco, 2006). This account does not require that attention fluctuates in sync with the stimulus, if attentional selection occurs at the level of the persistent perceived surface, rather than at the level of each alternate stimulus frame.

### Effect of alternation rate on attentional shifts in appearance

Across all 3 experiments we saw attention-based changes in appearance for stimuli alternating at 7.5Hz, which evoke a strong sense of transparency, but we also evidence of shifts in appearance for stimuli alternating at a slower rate (3.75Hz), which evokes a weaker sense of transparency (Vigano et al., 2015). This demonstrates that shifts in appearance of alternating stimuli are not limited to temporally transparent stimuli but are present where successive alternations are perceptually resolved. For stimuli alternating at 3.75Hz, our findings were mixed, with larger shifts than at 7.5Hz for simultaneously presented stimuli (Experiment 2), but no evidence of a shift in appearance when the stimuli were instead presented at the same location, across two intervals. These mixed findings may indicate an attentional process that is qualitatively different for 3.75Hz than at 7.5Hz. At both these alternation rates, changes in appearance could have been driven by attentional fluctuations in sync with the stimulus, while attentional selection of the persistent surface perception is likely to be stronger at 7.5Hz and possibly absent at 3.75Hz. It is unclear why attentional fluctuations that are in sync with the stimulus might operate differently for stimuli presented side by side versus across two intervals, but two factors are potentially relevant here. First, the side-by-side presentation could result in covert spatial attention being different between the two stimuli. Although the sides of the reference and target stimuli were counterbalanced across trials, since the reference stimulus contained an object silhouette or letter(s), it is likely that covert attention would be directed to the reference stimulus to a greater extent. Second, we deliberately randomised the starting phases of each stimulus to avoid synchronous phases providing a cue to differences in their stimulus cycles when they were presented side-by-side. This means that any entraining of attentional fluctuations in Experiments 1-2 likely occurred for one stimulus at a time, and was more likely to occur for the reference stimulus, whereas in Experiment 3 any attentional entrainment would have been more even for the two stimuli presented across two intervals. Although further work is necessary to test this account, if supported, this suggests that the shift in appearance for temporally transparent (7.5Hz) stimuli in Experiment 3 may reflect attentional selection of the persistent surface percept, which is much weaker at 3.75Hz.

### Feature binding in temporally transparent stimuli

Our findings also shed light on the phenomenon of temporal transparency, and the apparent paradox of feature binding for rapidly alternating stimuli (Holcombe & Cavanagh, 2001; Vigano et al., 2015).

In temporally transparent stimuli, (e.g. 7.5Hz alternation), each stimulus frame is only present very briefly (e.g. 67ms) before it is replaced by the alternate frame. Feature binding, such as perceiving the pairing of object color and orientation, is a relatively slow process with low temporal resolution (Fujisaki & Nishida, 2010; Holcombe, 2009; Treisman, 1996), and the brief duration of a single frame would be outside the range that feature binding can be achieved. Vigano et al. (2015) proposed that feature binding for these stimuli could be mediated by attentional selection of the contents of one frame: if the visual system represents the stimulus as two persistent, overlapping surfaces, then attentional selection of one surface could selectively boost the representation of the attended feature values and allow the readout of feature conjunction information. Our data support this account, since shifts in appearance of the more salient and task-relevant surface imply a selective enhancement of its neural representation, which would also facilitate the readout of feature conjunctions.

## Conclusion

We examined the effects of attention on the appearance of alternating stimuli and found that when stimulus and task conditions encouraged attention, the perceived dominance of the more salient and task-relevant stimulus increased. We found larger effects of salience, which increase the likelihood of exogenous attentional capture, but appearance was also shifted when only endogenous attention was engaged. Our results suggest that attention changed the appearance of these alternating stimuli, both in the temporally transparent range, and at slower alternation rates outside of this range. Shifts in appearance for temporally transparent stimuli provide evidence supporting the idea that attentional selection could mediate feature binding in temporal transparency.

## Acknowledgements

This project was funded by the Australian Research Council (ARC) (DP220100747 to EG). We thank Colin Clifford for helpful discussions on the manuscript.

## Declarations

### Ethics approval

The study was conducted according to the protocols approved by the Human Research Ethics Approval Panel – C (HREAPC) within the School of Psychology, UNSW (Reference iRECS4177).

### Consent to participate

Informed consent was obtained from all individual participants included in the study.

### Consent for publication

Patients signed informed consent regarding publishing their de-identified data.

### Availability of data and materials

The data and materials for both experiments are available in an OSF project that will be made public (URL added to manuscript) if accepted for publication.

### Code availability

Our manuscript includes all details necessary to recreate our experiment and analyses.

### Authors contributions

Conceptualization (ANI and EG); Investigation (ANI); Formal Analysis (ANI and EG); Writing – original draft (ANI and EG); Writing – review and editing (ANI and EG).

